# Rho/ROCK-dependent actomyosin contractility drives extracellular vesicle release from the cilium

**DOI:** 10.64898/2026.08.24.746798

**Authors:** William J. Spencer, Margaux J. Kreitman, Nicholas F. Schneider, Christin Hanke-Gogokhia, Stella Finkelstein, David G. Ball, Petar R. Mitev, Gregory J. Pazour, Vadim Y. Arshavsky

## Abstract

The release of extracellular vesicles (EV) from the primary cilium is a conserved process observed in many cell types. It serves as a rapid and efficient mechanism to release select proteins from the cilium, which can be used for either intercellular communication or membrane material disposal. Previous studies have shown that the release of EVs from the cilium relies on the actin cytoskeleton and proposed several molecular mechanisms that may perform this function. Using the model of IMCD3 cells, we now demonstrate that this process relies on actomyosin contractility supported by non-muscle myosin IIA acting downstream of the RhoA-ROCK signaling pathway. We further showed that the cilia of these cells release EVs independently of *de novo* actin polymerization, which we confirmed using an *in vivo* model of mutant photoreceptor cells that release massive amounts of vesicles from their cilia instead of elaborating into light-sensitive outer segment membrane structures.

## INTRODUCTION

The primary cilium is a sensory organelle present in many eukaryotic cells, which receives and processes a variety of external signals (see (Mill et al., 2023) for review). To modulate signaling activity, the primary cilium must tightly control its protein composition using specialized protein machinery, such as the IFT and BBSome particles, to selectively transport cargo into and out of the cilium. Defects in this machinery result in ciliary dysfunctions and cause syndromic diseases in humans, called ciliopathies, which are marked by severe impairments in the development and function of the retina, kidney, brain and other organs (Goetz and Anderson, 2010).

Another conserved mechanism which regulates the molecular composition of the cilium is the release of extracellular vesicles directly from the ciliary membrane (Luxmi and King, 2024; Luxmi et al., 2022; Ojeda Naharros and Nachury, 2022; Vinay and Belleannee, 2022; Wang and Barr, 2018). Because these vesicles are formed via an outward membrane budding and scission process they are categorized as *ectosomes* (aka *microvesicles*) (Wood et al., 2013). Ciliary ectosomes have been ascribed many functional roles, including protein disposal (Nager et al., 2017; Zhu et al., 2026), cell cycle reentry control (Phua et al., 2017), secretion of bioactive proteins (Luxmi et al., 2022; Walsh et al., 2022; Wood et al., 2013) and cell-to-cell communication (Wang et al., 2014).

Previous studies showed that ciliary ectosome release requires the action of actin filaments inside the cilium (Loukil et al., 2021; Nager et al., 2017; Phua et al., 2017; Prasai et al., 2024; Wang et al., 2019). An elegant experiment by Inoue and colleagues demonstrated that actin residing in the cilium is responsible for ectosome release by targeting a protein called thymosin beta-4 to the cilium that disrupts actin filaments and showed that this stopped ciliary ectosome release (Phua et al., 2017). However, there is no consensus about how actin in the cilium drives ectosome release. This process was reported to be sensitive to inhibitors of actomyosin contractility (Nager et al., 2017; Phua et al., 2017) and branched actin network expansion (Nager et al., 2017), as well as being regulated by MYO6 (Nager et al., 2017), drebrin (Nager et al., 2017), ACTN4 (Nager et al., 2017), INPP5E (Phua et al., 2017), Rab7 (Wang et al., 2019), CAPZB (Loukil et al., 2021) and CDC42 (Prasai et al., 2024).

In the present study, we revisited the mechanism by which polymerized actin regulates the release of ciliary ectosomes using the model of ciliated inner medullary collecting duct (IMCD3) cells. Our experiments demonstrate that ciliary ectosome release from these cells is strongly suppressed by inhibition of non-muscle myosin II isoforms or knockout of non-muscle myosin IIa but is unaffected by inhibition of branched actin polymerization. We further showed that the actomyosin contractility underlying ectosome release in these cells is dependent primarily on the myosin IIA isoform, which is regulated by an upstream RhoA-ROCK signaling pathway.

Our finding that ciliary ectosome release occurs independently from branched actin polymerization was further confirmed in an alternative, *in vivo* model of vertebrate photoreceptors. A unique property of the photoreceptor cell is that it builds an elaborate outer segment structure at its primary cilium, which serves to produce electrical responses upon capturing light (Goldberg et al., 2016; Spencer et al., 2020; Wensel et al., 2021). Outer segments consist of a stack of disc-shaped membranes, each formed upon an expansion of a branched actin network in a mechanism akin to lamellipodium formation in motile cells (Spencer et al., 2019; Spencer et al., 2023). Remarkably, a knockout of a protein called peripherin-2 causes the photoreceptor cilium to release a massive amount of ectosomes instead of making discs (Cohen, 1983; Salinas et al., 2017). We found that the knockout of WASF3 – the actin nucleation promoting factor indispensable for building the branched actin network at the site of photoreceptor disc formation – has no effect on ectosome release in this model, consistent with inhibition of branched actin polymerization in IMCD3 cells not affecting ectosome release.

## RESULTS

### Direct monitoring of ciliary ectosome release from IMCD3 cells

To directly monitor ciliary ectosome release in real time we conducted live-cell imaging experiments on IMCD3 cells stably expressing the fluorescently labeled ciliary marker, SSTR3. Cells were grown to confluence and subjected to serum starvation to induce ciliation. Ectosome release was measured by live-cell imaging the cilia from at least 100 cells for two hours with intervals between each frame varying from 30 seconds to 6 minutes. To ensure that entire cilia could be imaged in 3D throughout the time window, large Z-stacks were taken at each frame and axial focus drift was corrected in real time.

To increase the rate of ectosome release by these cells, we followed the strategy described in (Nager et al., 2017) and suppressed retrograde protein transport from the cilia by knocking out β-arrestin 2 (ARRB2). This increased the frequency of ectosome release by several-fold (**Fig. 1A**). At the basal state, ∼20% of ciliated *Arrb2^−/−^* cells released at least one ectosome from the ciliary tip during a two-hour timeframe (**Fig. 1A,B; Supplementary video 1**). This basal release rate was nearly doubled by supplementing the culture medium with 10% fetal bovine serum (FBS) at the start of imaging (**Fig. 1A**). We therefore added serum in all subsequent experiments. We also noted that ∼5% of cilia were completely shed by fragmentation, a type of deciliation that was previously shown to rely on the microtubule severing protein, katanin (Mirvis et al., 2019). These events were not counted as ectosome release events.

**Figure 1.**
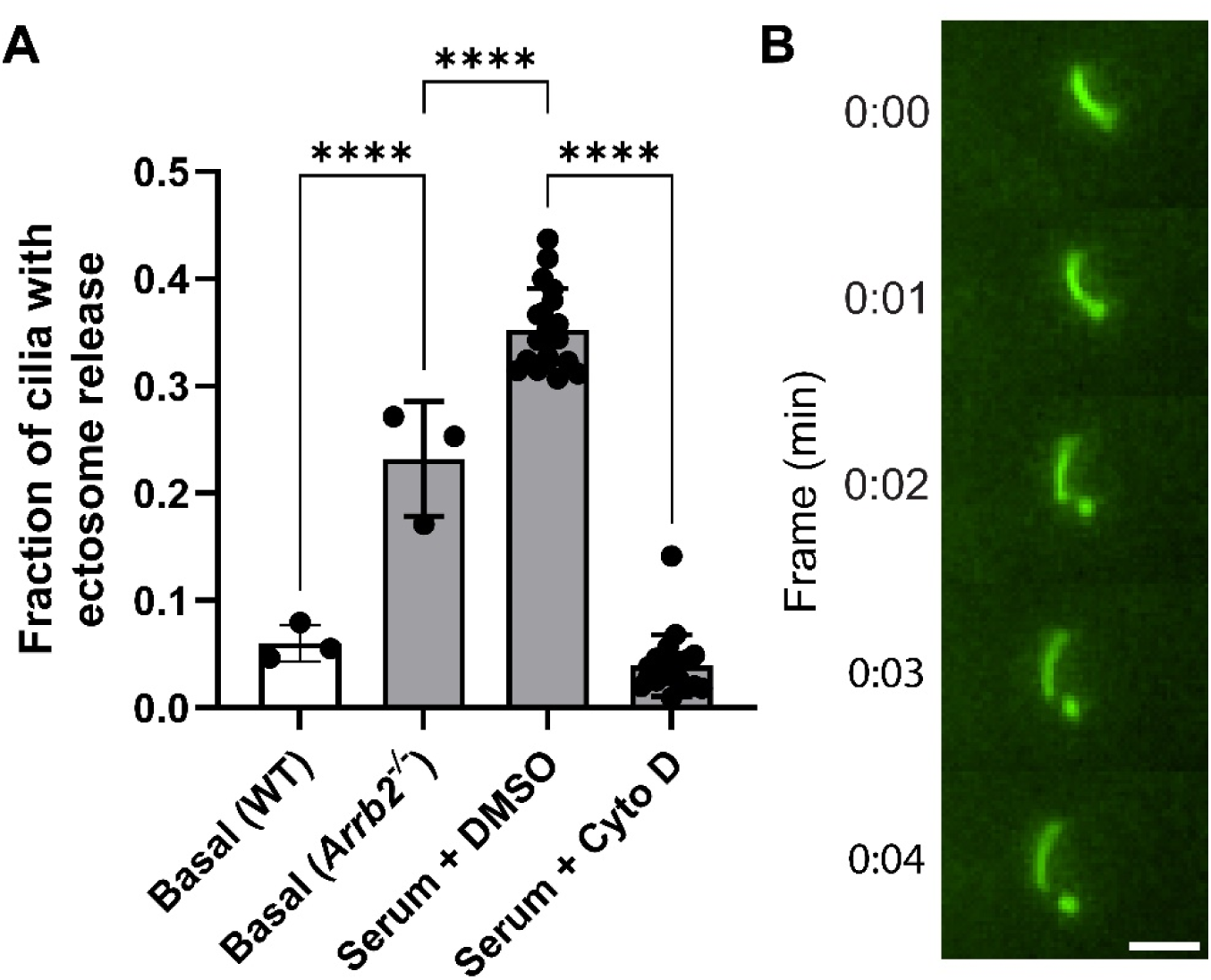
Ciliary ectosome release from IMCD3 cells is stimulated by serum and inhibited by cytochalasin D (Cyto D). **(A)** Quantification of ectosome release from the cilia of IMCD3 cells expressing SSTR3-EGFP, either WT (white) or *Arrb2^−/−^* (grey). Each data point represents an individual experiment conducted by live-cell imaging the cilia of at least 100 different IMCD3 cells. Statistical significance was assessed using one-way ANOVA. Data are shown as mean ± SD. Statistical annotation: **** p < 0.0001. **(B)** An example time lapse image series of an IMCD3 cell cilia releasing an ectosome. Scale bar is 2 µm.

As previously reported (Nager et al., 2017), ectosome release was almost completely suppressed by cytochalasin D added 1 hour before introduction of serum (**Fig. 2A**). Cytochalasin D binds to barbed ends of actin filaments thus preventing their further elongation. Because actin filaments are dynamic structures having constant monomer addition to barbed ends and monomer removal from the pointed end, cytochalasin D treatment leads to complete actin filament disassembly, which occurs on the timescale from a few minutes to hours (Goode et al., 2023). The dependency of ciliary ectosome release on cytochalasin D suggests the dynamic nature of actin filaments involved in ectosome release. In subsequent experiments we discerned amongst actin-dependent mechanisms potentially underlying this process.

**Figure 2.**
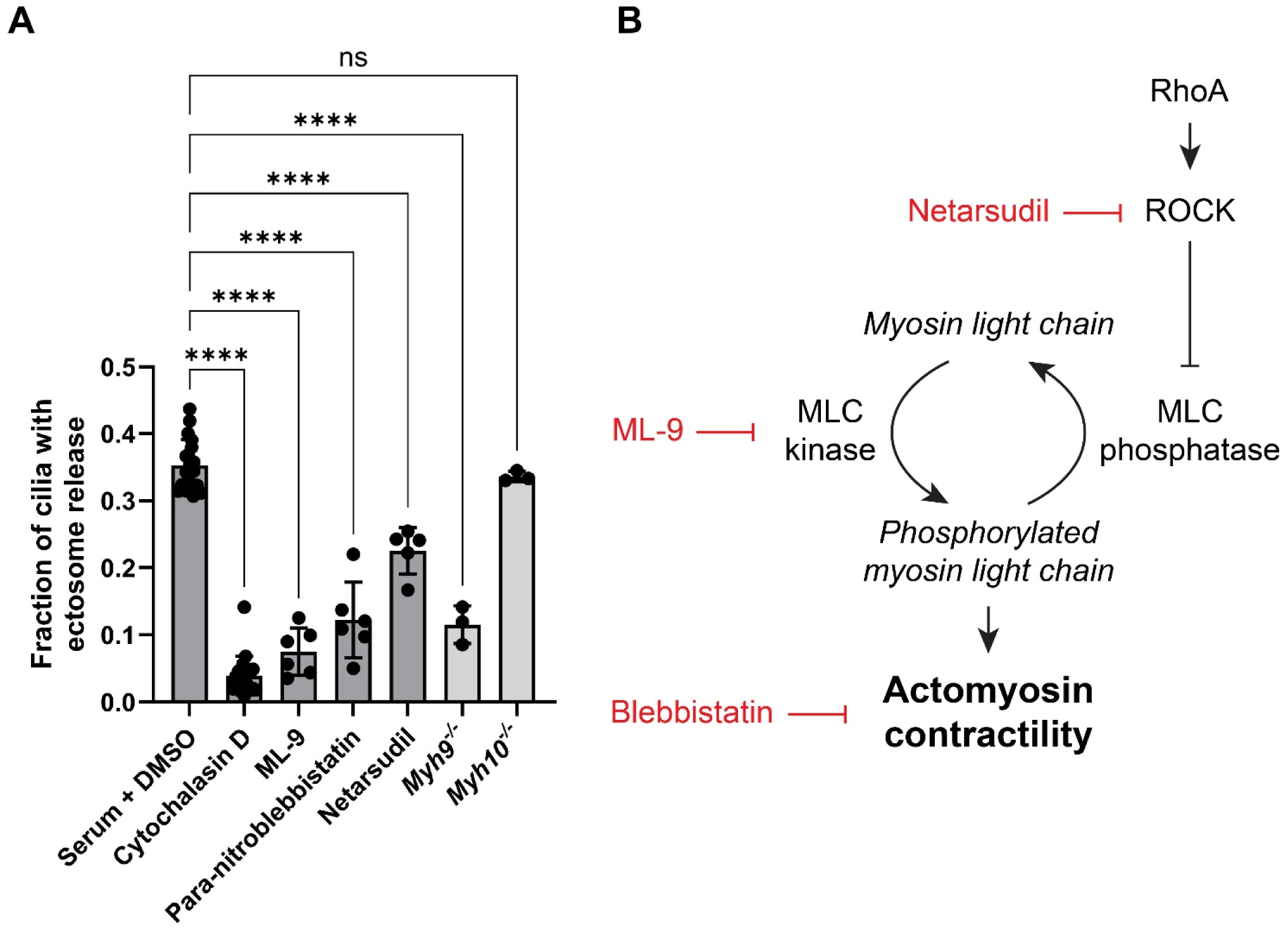
Inhibition of actomyosin contractility and upstream RhoA/ROCK signaling pathway suppress ciliary ectosome release. **(A)** Quantification of ectosome release from *Arrb2^−/−^* IMCD3 cells incubated with indicated drugs (at concentrations described in Methods) and knockout cells lacking non-muscle myosin IIa (*Myh9^−/−^*) or non-muscle myosin IIb (*Myh10^−/−^*). Each data point represents an individual experiment conducted by live-cell imaging of cilia from at least 100 different IMCD3 cells. Statistical significance was assessed using one-way ANOVA. Data are shown as mean ± SD. Statistical annotation: **** p < 0.0001. **(B)** A schematic of the RhoA/ROCK-dependent actomyosin contractility pathway with the drugs used in this study highlighted in red.

### Ciliary ectosome release from IMCD3 cells is inhibited by suppressing actomyosin contractility

In the next set of experiments, we took a pharmacological approach to elucidate the underlying actin-dependent mechanism. We first investigated the role of actomyosin contractility, which plays a crucial role in driving the membrane protrusion and release of membrane blebs. Blebs, like ectosomes, are outward protrusions of the plasma membrane that may be released from the cell (Fackler and Grosse, 2008; Wickman et al., 2013). Bleb formation requires non-muscle myosin II motor proteins to exert a constricting force on cortical actin that increases local hydrostatic pressure to produce a plasma membrane protrusion (Charras et al., 2008; Charras et al., 2005; Wickman et al., 2013). Consistent with previous reports (Nager et al., 2017; Phua et al., 2017), we found that ectosome release was markedly suppressed by para-nitroblebbistatin, a drug that keeps myosin in an actin-detached state and prevents actomyosin cross-linking (Kovacs et al., 2004) (**Fig. 2A**).

The activity of non-muscle myosin II is commonly regulated by the RhoA/ROCK pathway schematically illustrated in **Fig. 2B** (see (Guan et al., 2023) for a review). Actomyosin contractility is increased through phosphorylation of myosin light chain (MLC) by myosin light chain kinase (MLCK) and decreased through its dephosphorylation by myosin light chain phosphatase (MLCP). This equilibrium is regulated via phosphorylation of MLCP by Rho-kinase (ROCK), which inhibits the MLCP activity. ROCK is, in turn, activated by the small GTPase RhoA. Ultimately, the activation of RhoA/ROCK shifts the pool of MLC molecules toward the phosphorylated state, thus promoting actomyosin contractility (Kimura et al., 1996). Consistent with this mechanism being involved in the regulation of ciliary ectosome release from IMCD3 cells, we found that this release is strongly suppressed by the MLCK inhibitor ML-9, as well as the ROCK inhibitor netarsudil (**Fig. 2A**). Notably, netarsudil was a weaker inhibitor than ML-9 and suppressed ectosome release to a level comparable with the basal rate observed without FBS (cf. **Fig. 1A**). This suggests that the RhoA/ROCK pathway accounts for the enhanced ectosome release accompanying serum stimulation of these cells.

We next investigated the identity of the myosin motor responsible for ciliary ectosome release from IMCD3 cells. Because para-nitroblebbistatin inhibits all myosin II isoforms (Limouze et al., 2004) and because IMCD3 cells express only the IIA and IIB RNA transcripts (Chan et al., 2018) (coded by the *Myh9* and *Myh10* genes, respectively; **Supplementary Fig. 1**), we used CRISPR knockouts of myosin IIA and IIB to determine which isoform controls ectosome release from these cells. Whereas the knockout of myosin IIA decreased ectosome release to the same degree as para-nitroblebbistatin, knockout of myosin IIB did not have any effect (**Fig. 2A**). These data suggest that the myosin motor responsible for ciliary ectosome release in IMCD3 cells is non-muscle myosin IIA.

### Ciliary ectosomes are released from IMCD3 cells independent of *de novo* actin network expansion

In another group of experiments, we explored an alternative hypothesis that ectosome release involves de novo actin network expansion. Should this be the case, a burst in actin filament assembly would coincide with ectosome release from the cilium. The presence of F-actin in primary cilia is well established (Chaitin et al., 1984; Kiesel et al., 2020; Wang et al., 2023) and has been shown to partition proteins into distinct and highly fluid nanodomains throughout the cilium (Lee et al., 2018). Recent studies have shown an increase in actin filament assembly near the scission point between the ciliary ectosome and the cilium (Phua et al., 2017; Prasai et al., 2024). To test whether this phenomenon is observed in our system, we tracked the time course of actin filament assembly in the cilium of IMCD3 cells in which actin filaments were labeled with actin probes, SiR-Actin (**Fig. 3A; Supplementary video 2**) or Lifeact (**Fig. 3B; Supplementary video 3**). As in prior studies (Phua et al., 2017; Prasai et al., 2024), we took time lapse image series at a fast rate (<2 min/frame) to capture rapid changes in actin dynamics. We performed these experiments in confocal mode and used IMARIS software to 3D-segment the cilia. This allowed us to reliably distinguish between the actin dynamics occurring inside the cilium from that anywhere else in the cell.

**Figure 3.**
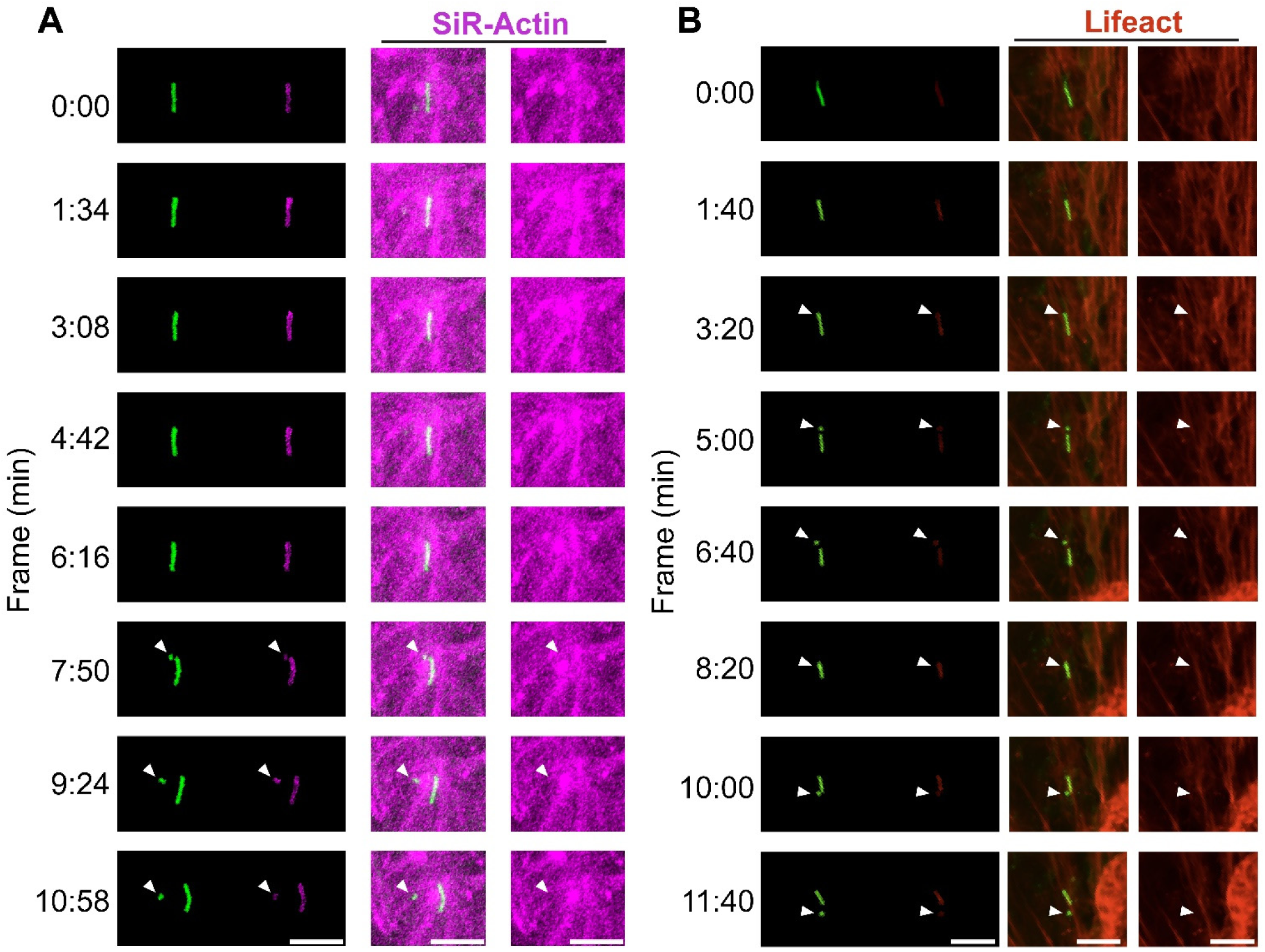
The abundance of intraciliary F-actin does not change with ciliary ectosome release in IMCD3 cells. **(A)** An example of a time lapse images series showing IMCD3 cell ciliary ectosome release. The cilium is labeled with SSTR3-EGFP and actin is labeled with SiR-Actin. The first column of images shows SSTR3-EGFP and F-actin channels after 3D segmentation of the cilium was performed in IMARIS. The two channels are separated from one another to specifically show the F-actin that is within the cilium. The middle column is a merged confocal image series without segmentation and the third column shows the F-actin staining alone. The white arrowhead marks the site where a ciliary ectosome is formed and released. Scale bar is 5 µm. **(B)** The same as in (A), but with actin labeled by Lifeact-mScarlet. Scale bar is 5 µm.

In contrast to previous reports, we did not detect any changes in intraciliary F-actin dynamics coinciding with or immediately preceding ectosome release (**Fig. 3**). Whereas we cannot explain this difference, it is worth pointing to multiple differences in experimental conditions, including both previous studies being conducted with mouse embryonic fibroblasts (Phua et al., 2017; Prasai et al., 2024). Further, a quantitative analysis in (Prasai et al., 2024) shows that the burst of F-actin at the site of ciliary ectosome release was observed only in a subset of cilia (at most ∼25%) and all illustrated examples show this burst to occur after rather than before ectosome release took place. Thus, it is conceivable that these observations reflect a tangentially related process involving *de novo* actin filament assembly. Regardless, our data indicate that, while relying on actomyosin contractility, ectosome release from IMCD3 cells does not require actin polymerization occurring concurrently with the release process.

### Branched actin is not required for ciliary ectosome release from vertebrate photoreceptor cells

A striking *in vivo* model to study ciliary ectosome release are photoreceptor cells from the peripherin-2 knockout mouse. Normal photoreceptors elaborate hundreds of disc-shaped membrane structures (or “discs”) stacked inside their cilium. This layered membrane arrangement serves to maximize light capture. However, the lack of peripherin-2 causes the photoreceptor cilium to release this membrane material in the form of ectosomes, which massively accumulate in the extracellular space surrounding these cells (Salinas et al., 2017). The accumulating vesicles were shown to be of ciliary origin (Salinas et al., 2017), and consistent with our findings in cell culture, they contain actin filaments (Chaitin et al., 1988), non-muscle myosin II and RhoA (Spencer et al., 2019).

The formation of each disc involves an evagination of the ciliary membrane at the base of the photoreceptor cilium (Burgoyne et al., 2015; Ding et al., 2015; Steinberg et al., 1980; Volland et al., 2015), and our previous study demonstrated that this evagination is driven by branched actin polymerization nucleated by Arp2/3 in a process akin the formation of lamellipodia in motile cells (Spencer et al., 2019). We further showed that the nucleation of this network requires the WAVE complex containing a unique WASF3 isoform and that this is the only Arp2/3 nucleation promoting factor detectable in the photoreceptor’s outer segment cilium (Spencer et al., 2023). Because this process occurs at the same ciliary location where ectosomes are released in the peripherin-2 knockout mouse, we decided to evaluate whether, in this case, branched actin contributes to ectosome release.

To test this hypothesis, we generated a double knockout mouse line lacking both peripherin-2 and WASF3 (*rds*^−/−^;*Wasf3^−/−^* mouse). We found that the knockout of WASF3 did not affect the massive release of ciliary ectosomes from photoreceptors of these animals (**Fig. 4**), essentially rejecting the role of branched actin in the process of ciliary ectosome release. We find this negative result particularly convincing because the genetic ablation of actin network formation at the photoreceptor cilium was constitutive and permanent.

**Figure 4.**
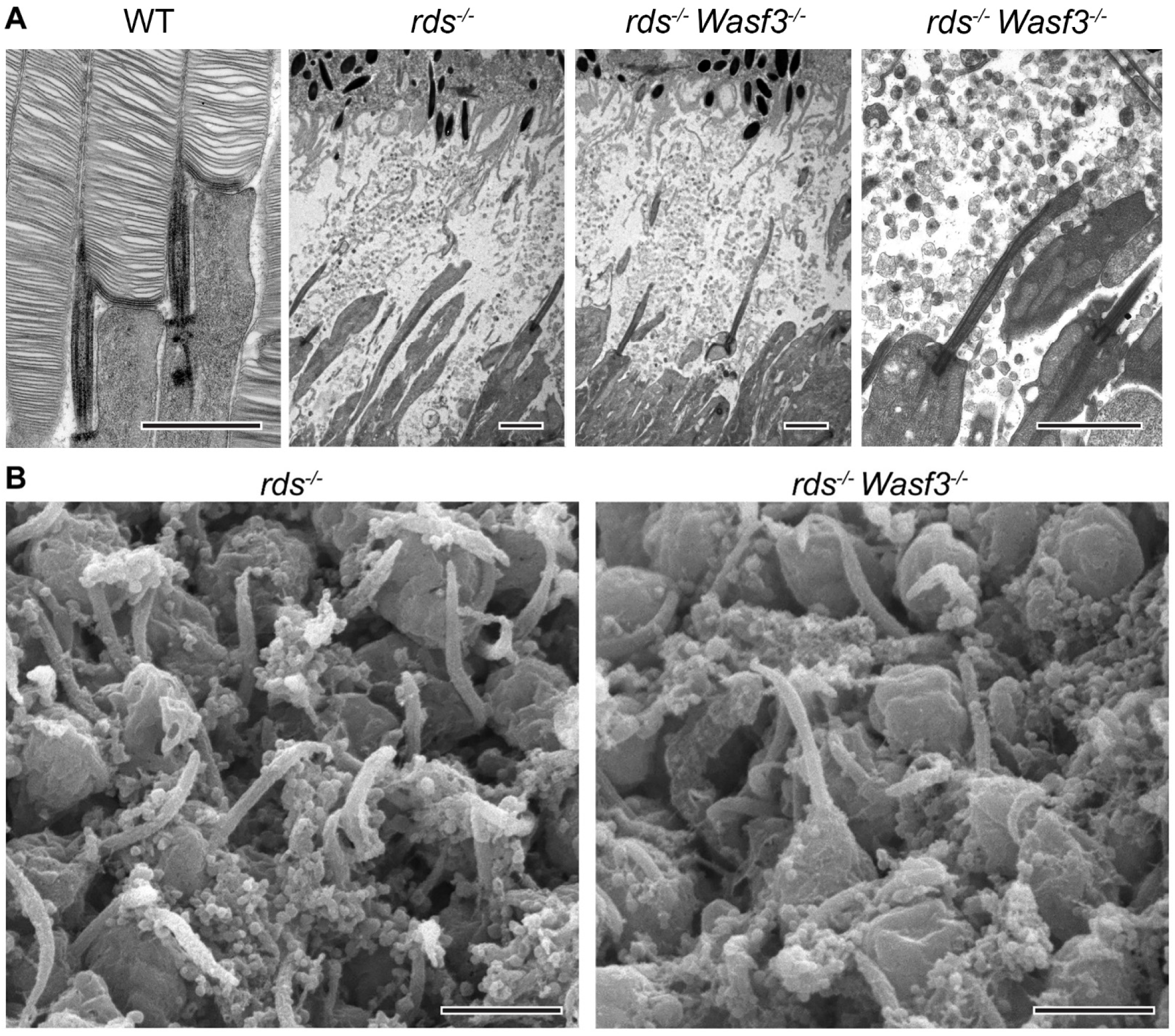
Absence of WASF3 has no effect on ectosome release in peripherin-2 knockout mice. **(A)** Transmission electron micrographs of rod outer segments from an adult WT mouse and of subretinal space from P14 *rds^−/−^* and *rds^−/−^ Wasf3^−/−^* mice. **(B)** Scanning electron micrographs of the surface of the retina from P14 *rds^−/−^*and *rds^−/−^ Wasf3^−/−^*mice. The photoreceptor side of the retina was imaged. Scale bars in all panels are 2 µm.

### Branched actin does not appear to be required for ciliary ectosome release from IMCD3 cells

In the final set of experiments, we additionally assessed the potential involvement of branched actin polymerization on ciliary ectosome release from IMCD3 cells by treating them with the Arp2/3 inhibitor CK-636, like in (Nager et al., 2017). The data shown in **Fig. 5** indicate that this treatment did not cause any effect on the rate of ciliary ectosome release when used at a concentration reported to efficiently suppress branched actin polymerization in cell culture (Oliveira et al., 2023; Sadhu et al., 2023; Zhu et al., 2023). This negative result appears to contradict a finding by Nager and colleagues (Nager et al., 2017) suggesting that CK-636 inhibits ectosome release in IMCD3 cells. A potential explanation for this difference is that the experiment in that study did not involve monitoring ectosome release but rather quantified the change in fluorescence intensity of the ciliary GPCR, NPY2R, presumably released from the cilium in the form of ectosomes. It is conceivable that this parameter may have been affected by CK-636 in a mechanism unrelated to ectosome release, for instance the dynamics of ciliary trafficking of NPY2R. Taking a lead from the mechanism of photoreceptor disc formation, we also tested whether the rate of ciliary ectosome release from IMCD3 cells is affected by the double knockout of Wasf1 and Wasf2, the two WAVE complex proteins expressed in these cells. Like in the case of Arp2/3 inhibition, this double knockout did not affect the rate of ectosome release (**Fig. 5**).

**Figure 5.**
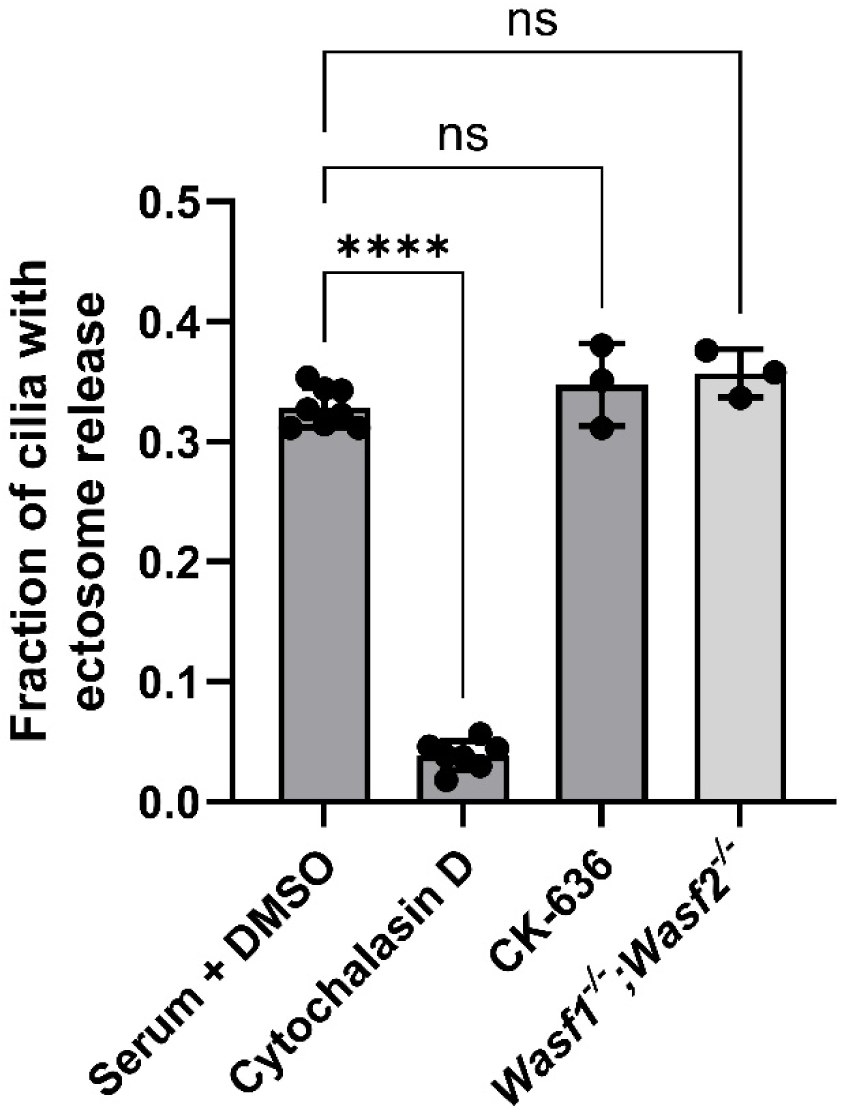
Inhibition of branched actin network does not affect ciliary ectosome release. Quantification of ectosome release from IMCD3 cells after: incubation with cytochalasin D; application of the Arp2/3 complex inhibitor CK-636; and double knockout of WASF1 and WASF2. Each data point represents an individual experiment conducted by live-cell imaging the cilia from at least 100 different IMCD3 cells. Statistical significance was assessed using one-way ANOVA. Data are shown as mean ± SD. Statistical annotation: *** p < 0.0001; ns = not significant.

## DISCUSSION

The central finding of this study is that actomyosin contractility involving non-muscle myosin IIa mediates ciliary ectosome release downstream of the RhoA/ROCK signaling pathway. We show that ciliary ectosome release is not accompanied by concurrent actin polymerization and does not rely on branched actin filaments. These properties suggest an analogy between the mechanisms underlying ciliary ectosome release and two related biological processes: blebbing and extracellular vesicle release from the plasma membrane.

In the case of blebbing, actomyosin contractility increases local hydrostatic pressure to push out a part of the plasma membrane (Charras et al., 2008; Charras et al., 2005; Wickman et al., 2013). The location of a bleb is thought to be determined by a disconnection of the linkages between cortical actin filaments and a specific segment of the plasma membrane, which allows this segment to be expanded upon an increase of pressure (Ikenouchi and Aoki, 2022). For example, it was shown that bleb formation can be induced by disconnecting cortical F-actin from ERM proteins, which link F-actin to the plasma membrane (Diz-Munoz et al., 2010; Fehon et al., 2010). The release of extracellular vesicles from the plasma membrane can also be driven by an actomyosin-mediated mechanism, as studied most extensively in a variety of cancer cells (Das et al., 2018; Li et al., 2012; Muralidharan-Chari et al., 2009; Schlienger et al., 2014; Sedgwick et al., 2015). The role of actomyosin contractility can extend to the formation of a contractile ring responsible for severing a vesicle from the plasma membrane, as observed in cytokinesis (Glotzer, 2005; Miller, 2011; Pollard and O’Shaughnessy, 2019). There are many reports that these processes are regulated by the RhoA/ROCK signaling pathway (Bement et al., 2005; Das et al., 2018; Li et al., 2012; Prokopenko et al., 1999; Schlienger et al., 2014; Sedgwick et al., 2015). Our data suggest that a similar molecular mechanism underlies ciliary ectosome release (**Fig. 6**). In this mechanism, RhoA activates ROCK, which phosphorylates and inhibits MLCP, resulting in the shift of non-muscle myosin IIA toward its active phosphorylated state. This leads to a localized constriction of the cilium resulting in release of an ectosome. Whether actomyosin contractility is sufficient to accomplish ectosome release or a concurrent detachment of membrane from the cortical F-actin inside the cilium is involved remains to be determined.

**Figure 6.**
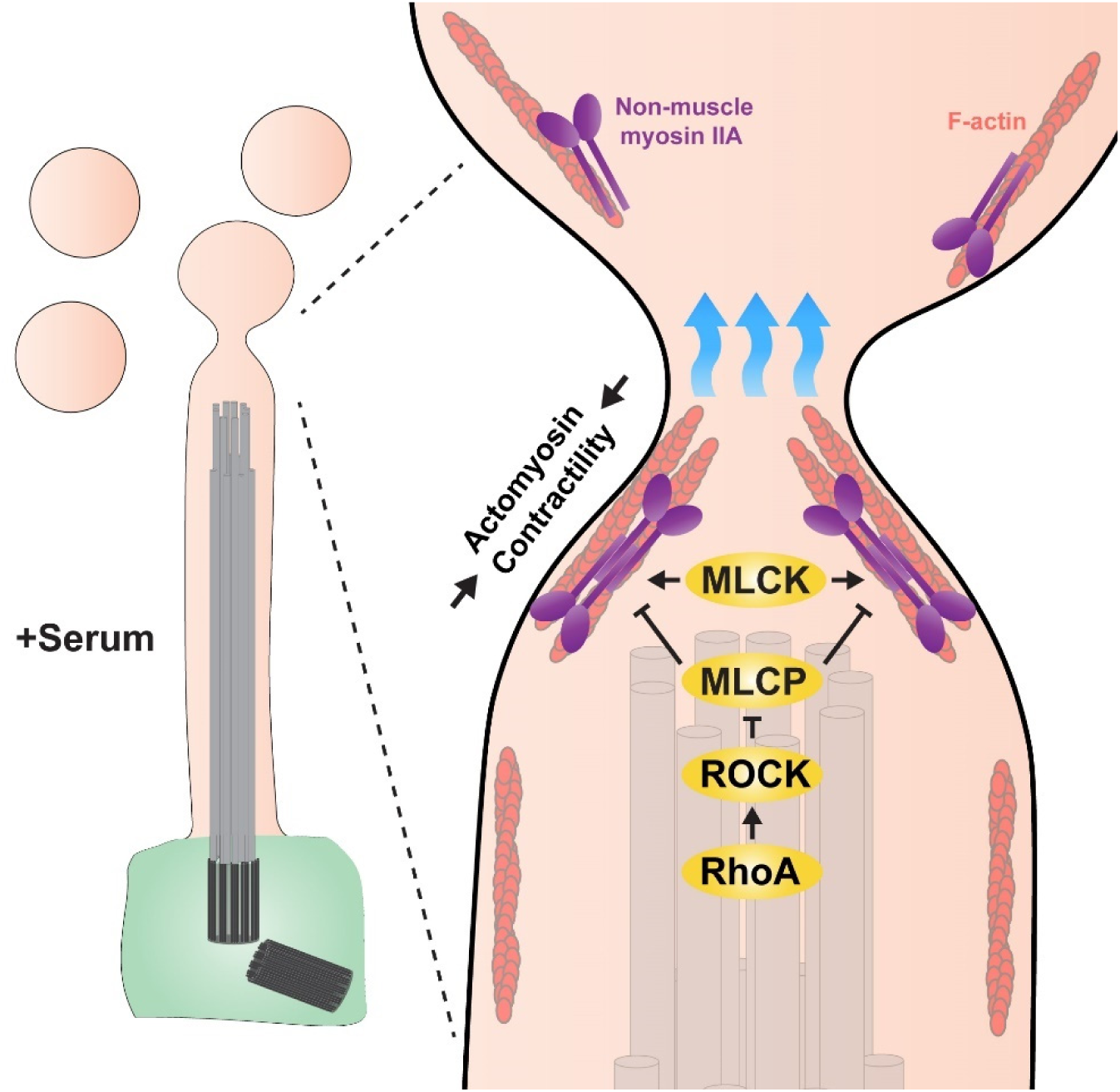
A model of ciliary ectosome release driven by RhoA/ROCK-modulated actomyosin contractility. In this model, intraciliary actomyosin contractility driven by non-muscle myosin IIA constricts the ciliary plasma membrane to push out an extracellular vesicle. This process is stimulated by MLCK and is regulated by the RhoA/ROCK signaling pathway targeting MLCP. RhoA, in its GTP bound state, activates ROCK, which phosphorylates and inhibits MLCP. Because MLCP normally suppresses actomyosin contractility, its inhibition shifts myosin to its active phosphorylated state, which induces contractility and facilitates the process of ectosome release.

Another interesting question arising from this study relates to the identity of the signal which could act upstream of the RhoA/ROCK pathway and ultimately trigger ectosome release. Such an upstream signal could originate directly in the cilium or elsewhere in the cell. For example, it has been shown that ectosomes could be released upon activation of the ciliary receptors SSTR3, NPY2R and smoothened (Cao et al., 2015; Nager et al., 2017; Zhu et al., 2026). On the other hand, the serum may initiate a signaling event elsewhere in the cell that is propagated to the cilium, presumably through the activation of RhoA. For example, serum contains lysophosphatidic acid (LPA), which is known to activate GPCRs coupled to G proteins G_12/13_ (Kranenburg et al., 1999; Siehler, 2009). The major established function of G_12/13_ is to stimulate RhoA guanine nucleotide exchange factor proteins (such as p115-RhoGEF) that catalyze nucleotide exchange on RhoA, shifting it to the active GTP-bound form (Siehler, 2009; Yu and Brown, 2015). Consistent with the involvement of LPA in ciliary ectosome release is the fact that it is the major factor in serum that drives ciliary disassembly (Hu et al., 2021) – a process reported to involve ectosome release from the cilium (Phua et al., 2017). More experiments are needed to elucidate how these signals communicate with the RhoA/ROCK actomyosin contractility pathway to drive ectosome release. In some cases of ciliary ectosome release an external signal may not be required, such as in the case of baseline release in our experiments. Instead, release appears to occur in response to accumulation of protein material in the cilium (Razzauti and Laurent, 2021; Salinas et al., 2017; Wang et al., 2021) or from mechanical stimulation (Wang et al., 2020).

A closely related question relates to biological functions performed by ectosomes following their release from the cilium. A study by Wang and colleagues showed that ciliary ectosomes released by *Caenorhabdidits elegans* induce male tail-chasing behaviors in other *C. elegans* animals, potentially increasing their reproductive success (Wang et al., 2014). Subsequent studies identified that ciliary ectosomes can act as vehicles to transfer proteins and RNA from one *C. elegans* animal to another during the mating process (Nikonorova et al., 2022; Wang et al., 2020). Ciliary ectosomes also aid in the mating of *Chlamydomonas* by delivering enzymes that facilitate the rupture of the sporangium wall (Wood et al., 2013) or serving as a chemoattractant to mates through peptidergic signaling (Luxmi and King, 2023; Luxmi et al., 2019). Other biological functions of ciliary ectosomes are cell autonomous. By rapidly discarding proteins via vesicles, ciliary ectosome release serves as a means to quickly alter the protein composition of the cilium. For example, Nager and colleagues showed that SSTR3 is packaged into ectosomes more efficiently when stimulated by its ligand and less efficiently when ciliary ectosome release is stimulated by a ligand for a different signaling pathway (Nager et al., 2017). Ciliary ectosome release was also shown to regulate cell cycle reentry, perhaps through enhancing hedgehog signaling (Phua et al., 2017). Recently, it was shown that the content of ciliary ectosomes can be heterogeneous and sub-populations of these vesicles can be specifically packaged with related signaling molecules, such as the polycystin proteins *lov-1* and *pkd-2* (Nikonorova et al., 2025; Wang et al., 2026). Although the role of polycystin-enriched ectosomes is unknown, they have been proposed to be “signalosomes” that play a role in *C*. *elegans* communication (Walsh et al., 2022) or, in humans, could play a role in the pathogenesis of polycystic kidney disease (Ding et al., 2021).

Finally, the results obtained in our study shed light on the understanding of evolutionary origins of photoreceptor disc morphogenesis. Whereas discs are built from the membrane material otherwise destined to be released from the cilium as ectosomes, their formation does not employ actomyosin contractility. Rather, they are shaped through a mechanistically distinct “lamellipodia-like” process involving the propagation of a branched actin network.

In conclusion, our study provides a mechanistic framework for understanding ectosome release from the primary cilium. The challenge of future experiments is to address several questions raised in this study, such as the precise mechanism of membrane scission or the versatility of cellular pathways controlling ectosome release.

## METHODS

### Animals

Mice handling was performed in accordance with the approved protocol by the Institutional Animal Care and Use Committees of Duke University. *Wasf3^−/−^* mice (Qin et al., 2019) were generously provided by Dr. John Cowell (Georgia Cancer Center) and crossed with *rds^−/−^* mice (stock no. 001979) obtained from Jackson Labs. All mice were housed under a 12/12-hour diurnal light cycle.

### Antibodies

We used rabbit polyclonal antibodies against non-muscle myosin IIA (3403; Cell Signaling Technology), rabbit polyclonal antibody against non-muscle myosin IIB (3404; Cell Signaling Technology), and rat monoclonal antibody to HSC70 (ADI-SPA-815; Enzo).

### Cell culture

The cell line used in this study was generated by lentiviral transfection of murine IMCD3 cells with GP814, which expresses SSTR3-EGFP from an EF1α promoter. Cells were cultured in the DMEM-F12 medium (02490; Gibco) supplemented with 10% FBS (F0926; Sigma), 100 U/ml penicillin and 100 ug/ml streptomycin (SV30010; Cytiva). Cells were occasionally checked for mycoplasma contamination using MycoGuard Mycoplasma PCR Detection Kit 2.0 (MP004; GeneCopoeia).

### CRISPR knockouts

*Arrb2* was knocked out from the SSTR3-EGFP expressing cells by CRISPR-Cas9 using two guide RNA sequences: 5’-GGTGCTGAAGAGGCAAATGT-3’ and 5’-GGGACCTCCGGTCAGACATG-3’ cloned into a PX458 vector (Addgene plasmid #48138), which contains Cas9 fused to mCherry. Dually EGFP and mCherry-positive cells were clonally selected by cell sorting. To knockout *Myh9*, *Myh10*, *Wasf1* and *Wasf2*, *Arrb2^−/−^* IMCD3, cells expressing SSTR3-EGFP were transiently transfected with the PX459 vector (Addgene plasmid #48139) modified by replacing the puromycin-resistant cassette with a neomycin resistance cassette. This vector contained one of the following sgRNA sequences: CAACGAAGCTTCGGTGCTGC for *Myh9*, TTGAAGGACCGCTACTATTC for *Myh10*, GGCTGAGCTCAAGATGCCGT for *Wasf1*, and GGAACATCGAGCCAAGGCAC for *Wasf2*. Knockout cells were selected with 80 µg/ml G-418 in FBS-supplemented DMEM:F12 media for 72 hr, allowed to expand in FBS-supplemented DMEM:F12 for 24 hr, and clonally selected by dilute plating in 96-well plates. Clones were assayed by PCR for mutations in the vicinity of the sgRNA sequences.

### Live cell imaging

IMCD3 cells expressing SSTR3-EGFP were grown to 90% confluence in FBS-supplemented DMEM:F12 media in coverslip-bottom 24-well plates (P24-1.5H-N; Cellvis). Ciliation was induced by removing FBS for 24 hours. Ectosome release was stimulated by reapplication of 10% FBS immediately prior to imaging. Plates were maintained at 37°C in humidified air containing 5% CO_2_ using an Okolab stage top incubator. For quantification of ectosome release, epifluorescence microscopy images of the cilia were acquired from at least 10 distinct locations in each of four to six wells using a 1.49 NA 60x objective on a Nikon Eclipse Ti2 microscope equipped with a Nikon DS-Qi2 camera and a motorized stage. At each imaging location, the microscope’s focal plane was adjusted to focus on the upward-facing cilia that can release ectosomes into the media. *Z*-stacks spanning 7 µm were acquired with 0.7 µm *z*-steps, and axial drift was corrected in real-time using Nikon Perfect Focus System. We programmed the microscope to image continuously for 2 hours intervals ranging from 30 s to 6 min. To quantify ciliary ectosome release, at least 100 cilia were analyzed under each condition and those drifted out of our observation volume were discarded. Image analysis and processing was performed with ImageJ. For imaging actin dynamics during ectosome release, we labeled F-actin in live-cells using SiR-actin (CY-SC001; Cytoskeleton Inc.) or by transfecting the cells with a plasmid expressing Lifeact fused to mScarlet (Bindels et al., 2017). To simultaneously image F-actin and SSTR3-EGFP in these experiments, we used the Nikon AXR resonant scanning confocal for dual channel image acquisition at ∼2 min intervals resulting in ∼60 images per location. The cilia in 3D confocal images were segmented in IMARIS software based on manually defined thresholds for SSTR3-EGFP fluorescence intensity and voxel number. The segmentation was applied across all imaging channels allowing visualization of actin staining within the segmented cilia.

### Drug treatments

The following drugs were used at indicated concentrations: 1 µM Netarsudil, 5 µM CK-636 (HY-15892; MedChemExpress), 10 µM cytochalasin D (C8273; Sigma), 10 µM N-paranitroblebbistatin (24171; Cayman) and 50 µM ML-9 hydrochloride (0431; Tocris). The ciliated IMCD3 cells were pre-incubated with Netarsudil for 4.5 hours, CK-636 for 2 hours, cytochalasin D for 1 hour, N-paranitroblebbistatin for 1 hour and ML-9 hydrochloride for 1 hour before addition of serum.

### Transmission electron microscopy

Fixation and processing of mouse eyes for thin plastic sections was performed as described previously (Ding et al., 2015). Mice were deeply anesthetized and transcardially perfused with 2% paraformaldehyde, 2% glutaraldehyde, and 0.05% calcium chloride in 50 mM MOPS (pH 7.4) resulting in exsanguination. The eyes were enucleated and fixed for an additional 2 hours in the same buffer at 22 °C. The fixed eyes were washed in PBS, eyecups were dissected and embedded in PBS containing 2.5% agarose (KU VF-AGT-VM; Precisionary), and cut into 200 µm thick slices on a vibratome (VT1200S; Leica). The vibratome sections were stained with 1% tannic acid (Electron Microscopy Sciences) and 1% uranyl acetate (Electron Microscopy Sciences), gradually dehydrated with ethanol and embedded in Spurr’s resin (Electron Microscopy Sciences). 70 nm sections were cut, placed on copper grids, and counterstained with 2% uranyl acetate and 3.5% lead citrate (19314; Ted Pella). The samples were imaged on a JEM-1400 electron microscope (JEOL) at 60 kV with a digital camera (Biosprint 16; AMT).

### Scanning electron microscopy

Mouse retinas were dissected in ice cold Ringer’s solution containing 130 mM NaCl, 3.6 mM KCl, 2.4 mM MgCl_2_, 1.2 mM CaCl_2_, and 10 mM Hepes (pH 7.4), adjusted to 314 mOsm. The retinas were fixed with 2% paraformaldehyde, 2% glutaraldehyde, and 0.05% calcium chloride in 50 mM MOPS (pH 7.4) for 1 hour with gentle agitation. Retinas were then washed with PBS and post-fixed with 1% Osmium Tetroxide in PBS (Electron Microscopy Sciences – EMS) for 1 hour. Next, the retinas were washed with PBS, subsequently dehydrated through graded ethanol to 100% and finally dried via the critical point method utilizing a Tousimis Samdri PVT-3D. Dried retinas were then mounted onto 12mm SEM stubs with carbon conductive paint (EMS) and immediately sputter coated with ∼15 nm of Au-Pd alloy utilizing a Denton Desk-V coater. Lastly, retinas were analyzed with a JEOL JSM-IT100 Scanning Electron Microscope operated at 10kV, and digital micrographs collected with an Everhart-Thornley secondary electron detector.

## ACKNOWLEDGEMENTS

This work was supported by the National Institutes of Health grants EY012859 (V.Y.A.), EY035525 (V.Y.A.), EY005722 (V.Y.A.), GM060992 (G.J.P.), T32EY037212 (D.G.B.), the Career-Starter Research Grant from the Knights Templar Eye Foundation (W.J.S.), Career Development Award from Research to Prevent Blindness (W.J.S), Career Development Award from The E. Matilda Ziegler Foundation for the Blind (W.J.S) and Unrestricted Awards from Research to Prevent Blindness Inc. (Duke University and SUNY Upstate Medical University).

## SUPPLEMENTARY MATERIALS

**Supplementary Fig. 1.**
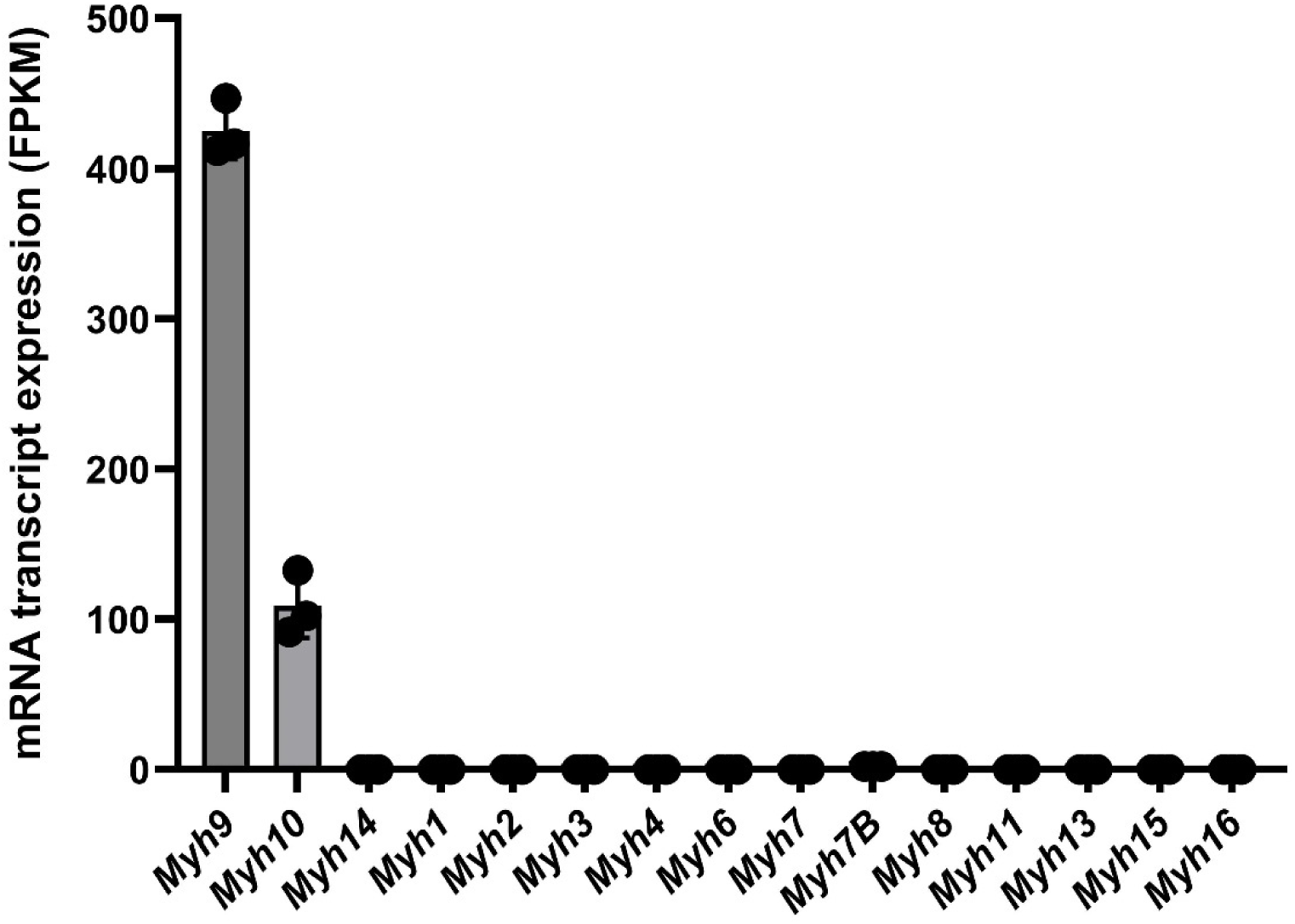
The mRNA expression levels of all blebbistatin-sensitive, class 2 myosin heavy chains in IMCD3 cells measured by RNA sequencing. The transcript levels are displayed as fragments per kilobase of transcript per million mapped reads (FPKM). The raw data are from (Chan et al., 2018).

**Supplementary video 1.** A time lapse video showing an example of ciliary ectosome release from a *Arrb2^−/−^* IMCD3 cell expressing SSTR3-EGFP. Sampling rate is 1 min. Scale is bar 2 µm.

**Supplementary video 2.** A time lapse video showing segmentation of ciliary ectosome release from a *Arrb2^−/−^* IMCD3 cell expressing SSTR3-EGFP. The SiR-Actin signal within the cilia was shifted to the right of the SSTR3-EGFP signal allowing simultaneous visualization of both channels. Sampling rate is 1 minute and 34 seconds. Scale bar is 1 µm.

**Supplementary video 3.** A time lapse video showing segmentation of ciliary ectosome release from a *Arrb2^−/−^* IMCD3 cell expressing SSTR3-EGFP. The Lifeact-mScarlet signal within the cilia was shifted to the right of the SSTR3-EGFP signal allowing simultaneous visualization of both channels. Sampling rate is 1 minute and 40 seconds. Scale bar is 1 µm.

## Abbreviations

EV: extracellular vesicle

## Notes

### Competing Interest Statement

The authors have declared no competing interest.

